# A multi-method approach for proteomic network inference in 11 human cancers

**DOI:** 10.1101/015214

**Authors:** Yasin Şenbabaoğlu, Selçuk Onur Sümer, Giovanni Ciriello, Nikolaus Schultz, Chris Sander

## Abstract

Protein expression and post-translational modification levels are tightly regulated in neoplastic cells to maintain cellular processes known as ‘cancer hallmarks’. The first Pan-Cancer initiative of The Cancer Genome Atlas (TCGA) Research Network has aggregated protein expression profiles for 3,467 patient samples from 11 tumor types using the antibody based reverse phase protein array (RPPA) technology. The resultant proteomic data can be utilized to computationally infer protein-protein interaction (PPI) networks and to study the commonalities and differences across tumor types. In this study, we compare the performance of 13 established network inference methods in their capacity to retrieve literature-curated pathway interactions from RPPA data. We observe that no single method has the best performance in all tumor types, but a group of six methods, including diverse techniques such as correlation, mutual information, and regression, consistently rank highly among the tested methods. A consensus network from this high-performing group reveals that signal transduction events involving receptor tyrosine kinases (RTKs), the RAS/MAPK pathway, and the PI3K/AKT/mTOR pathway, as well as innate and adaptive immunity signaling, are the most significant PPIs shared across all tumor types. Our results illustrate the utility of the RPPA platform as a tool to study proteomic networks in cancer.

**Availability:** PPI networks from the TCGA or user-provided data can be visualized with the **ProtNet** web application at http://www.sanderlab.org/protnet/.

## INTRODUCTION

### The utility of high-throughput proteomic datasets for probing cancer-related pathways

The Cancer Genome Atlas (TCGA) Research Network has recently profiled and analyzed large numbers of human tumors both within and across tumor lineages to elucidate the landscape of cancer associated alterations at the DNA, RNA, protein, and epigenetic levels ^1^. Integrated analyses of the resulting rich genetic and epigenetic data types have already started to shed light on commonalities, differences and emergent themes across tumor lineages ^2, 3^. Analysis of TCGA samples from 11 tumor types indicated that whole protein and phosphoprotein levels in these tumors, as measured by antibodies on reverse phase protein arrays (RPPA), capture information not available through analysis of DNA and RNA ^4^. The RPPA platform used in the Akbani *et al*. analysis was subsequently expanded to include 187 high quality antibodies targeting 136 proteins and 51 phosphoproteins, i.e. phosphorylated states of proteins. These antibodies were selected with a focus on cancer-related pathway and signaling events and analyzed with the intent to discover new therapeutic opportunities. This dataset is available for download from The Cancer Proteome Atlas ^5^, and referred to as PANCAN11 from here on.

### Analysis of function requires knowledge of interactions

The availability of proteomic datasets such as PANCAN11 where protein levels are measured across different conditions pose a unique opportunity to study the functions of proteins. However, the analysis of function requires knowledge of interactions. For instance, in the protein-folding domain, the function of a single residue during folding can be determined only by having knowledge about the residues it is interacting with. Similarly, the functions of a protein in the cell can only be understood by determining the interaction partners. Therefore, the units of analysis are not the individual protein expression levels, but the interactions of proteins with other cellular entities.

Statistical tools such as correlation can be used to study the interactions of proteins. However, correlation between two proteins does not imply that they **directly interact**, because correlation may also be induced by chaining of correlation between a set of intervening, directly interacting proteins. Such indirect correlations are called **transitive interactions**. It was previously shown that the dominant correlations in a system can be the result of parallel transitive interactions ^6^.

There are three main network motifs that lead to transitive interactions: fan-in, fan-out and cascade. A **fan-in** is a case where there are direct interactions from proteins A and B to a third protein C but there is no interaction between A and B. A **fan-out** is the situation where there is a direct interaction from protein C to both A and B but there is no interaction between A and B. A **cascade**, on the other hand, is a chain event where there are direct interactions from A to B, and from B to C, but not from A to C. In all these three cases, if the two direct interactions are in the same direction (both positive or both negative), there is a transitive influence observed between the proteins that do not have a direct interaction. Since biological pathways and signaling events contain many fan-in, fan-out and cascade network motifs, transitive effects occur widely across the network and have previously been shown to be a systematic source of false positive errors for many computational network inference methods ^7^. Thus, it is crucial to minimize the effect of transitive interactions in building network models from high-throughput datasets.

### A diverse array of computational network inference methods

A wide array of computational methods has been proposed in the literature for the identification of direct interactions in networks. The common objective of many of these methods is to call a direct interaction between two entities if they are *‘not conditionally independent’* of each other given a set of other entities. One simple example is the regression-based **partial correlation** approach. Consider a three-variable system consisting of A, B, and C. When testing the existence of a direct interaction between A and B in this approach, measurements on A and B would first separately be regressed on the measurements on C, the residual vectors would be computed, and then the correlation between the residual vectors would be found. If this *‘partial’* correlation is significantly different from zero, a direct interaction is called between A and B.

Despite the similarity in the objective, these methods employ diverse inference procedures such as **mutual information** ^8-11^, **regression** ^12-14^, **Gaussian graphical models** ^15, 16^, and **entropy maximization** ^17, 18^. The diversity of algorithms for inferring direct interactions, coupled with the absence of a robust off-the-shelf method, creates challenges for users that aim to generate hypotheses and eventually discover novel functional interactions among proteins. We address this challenge by testing different families of network inference methods towards the goal of deriving guidance for the better-performing methods.

### Evaluating the performance of network inference methods on a pan-cancer proteomic dataset

The RPPA platform, first introduced in Paweletz *et al.* ^19^ stands a good chance of becoming a widely used proteomics platform as greater numbers of reliable antibodies are being developed. Here, we present a rigorous comparison of the performance of 13 commonly used network inference algorithms based on PANCAN11, a pan-cancer RPPA dataset, which contains levels of many (phospho)proteins in a large number of samples, such that reasonably meaningful protein-protein correlations can be computed. The goal of this comparison is two-fold: To investigate 1) if the signal-to-noise ratio of the RPPA technology allows the discovery of known and novel PPIs, 2) to what extent algorithms that were originally developed for gene regulatory network inference accomplish the inference of PPIs.

Performance evaluation of PPI network inference for different cancers requires a ‘gold-standard’ for each cancer type. However, a true gold-standard for human PPIs does not exist, let alone a separate one for each tumor type. Most protein interactions in *in vivo* systems remain unknown or unproven and/or depend on physiological context. Yet public knowledgebases that store collections of curated pathway and/or interaction data contain useful information. For instance, **Pathway Commons (PC)** is a collection of publicly available and curated physical interactions and pathway data including biochemical reactions, complex assembly, transport and catalysis events ^20^, aggregated from primary sources such as Reactome, KEGG and HPRD and conveniently represented in the BioPAX pathway knowledge representation framework ^21-24^.

In this study, we adopted PC as a benchmark, and evaluated the performance of 13 network inference methods in their capacity to retrieve ‘true’ protein-protein interactions from RPPA datasets of 11 cancer types. We then used a group of high-performing methods to investigate the similarities and differences among the 11 cancer types in our dataset. The workflow of this study (**Figure 1**), involves the parallel generation of two PPI network models, one from *computational inference* and one from the *pathway knowledgebase*. On the inference side, multiple antibodies are assayed on an RPPA platform (Step 1) and the resulting dataset is normalized to generate a proteomic profile of the cohort such as PANCAN11. Computational network inference methods are then employed to create a network model with the inferred PPIs (Step 2). On the knowledgebase side, various wet-lab experiments are performed to generate data, and the resulting information is stored in the scientific literature (Step 3). Curators sift through the literature to distill multiple-layered information on PPIs (Step 4), and then this information is catalogued in knowledgebases such as PC (Step 5). A comparison of the PPI network models from the two sides reveals the level of fidelity at which the ‘true’ network is constructed by the computational methods (Step 6).

**Figure 1:**
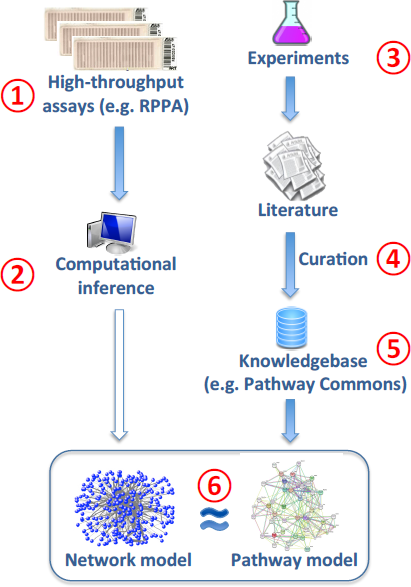
Workflow for the performance evaluation of network inference methods on a proteomic dataset. The workflow is comprised of a computational inference component and a pathway knowledgebase component that are used to generate separate PPI network models. Caveats involved in Steps 1-6 are discussed in **Discussion** and **Text S1**.

### Ascertainment bias in Pathway Commons

The workflow of performance evaluation as described above involve certain caveats. These are discussed in detail in the **Discussion** section and in **Text S1**. Here we discuss one of the caveats, the *ascertainment bias* in pathway knowledgebases (Step 5 in **Figure 1**). Wet-lab experiments for PPI plausibly have over-representation of certain proteins due to the perceived interest in the field and ease of study. In a recent paper, a Pearson correlation of 0.77 was reported for the correlation between the number of publications in which a protein was mentioned and the number of interactions reported for that protein in literature-curated data ^25^. This implies the potential existence of an ascertainment bias in pathway knowledgebases. There will exist more documented interactions of a certain protein if that protein is studied more intensively by the community. The *ascertainment bias* in PC precludes our benchmark network from being a *true gold standard*. However, in the absence of a true gold standard, we adopted this network as a *working gold standard*. This and other caveats challenge the comparability of pathway models from a knowledgebase and network models from a computational algorithm. Thus, it is necessary to be mindful of these caveats when interpreting the performance evaluation results in this study.

## RESULTS

We evaluated in this study the performance of 13 different network inference methods on the PANCAN11 RPPA dataset by using PC as a benchmark. The PANCAN11 dataset is comprised of 3467 samples and 187 proteins. The total number of possible non-self interactions with 187 proteins is 17,391. However, the number of interactions in PC (version 2) involving any two proteins from this set of 187 is 1,212 as determined by *PERA* ^26^. This PC benchmark subnetwork of 1,212 interactions forms the working gold-standard for this study, and has only 162 of the 187 proteins as interaction partners (meaning PC does not have any interactions between the remaining 25 proteins and any one of the full set of 187 proteins). As 162 proteins can form a total of 13,041 non-self interactions, the gold standard network has a density of 9.29%.

### *Limited-recall* versus *full-recall* for the optimization of precision-recall curves

We obtained network predictions for 11 tumor types listed in **Table 1** by using the 13 network inference methods listed in **Table 2**. We employed the **precision – recall** curves to first find the optimal parameter values for each method, and then to compare the performance of methods using their optimal values. The precision-recall (PR) curves were constructed by first ranking an edge list based on significance, and then plotting precision and recall on the **y** and **x** axis respectively for cumulatively increasing numbers of the top (the most significant) edges from the list. The trade-off between precision and recall at different cutoffs gives a reliable idea about the performance of a method, and this performance can be quantified with the area under the precision-recall curve (AUPR).

**Table 1:** PANCAN11 tumor types, the abbreviations used in the study, and the number of samples in each tumor type

| **Tumor type** | **Abbreviation** | **Number of samples** |
| --- | --- | --- |
| Bladder urothelial carcinoma | BLCA | 127 |
| Breast invasive carcinoma | BRCA | 747 |
| Colon adenocarcinoma | COAD | 334 |
| Glioblastoma multiforme | GBM | 215 |
| Head and neck squamous cell carcinoma | HNSC | 212 |
| Kidney renal clear cell carcinoma | KIRC | 454 |
| Lung adenocarcinoma | LUAD | 237 |
| Lung squamous cell carcinoma | LUSC | 195 |
| Ovarian serous cystadenocarcinoma | OVCA | 412 |
| Rectum adenocarcinoma | READ | 130 |
| Uterine corpus endometrioid carcinoma | UCEC | 404 |
| **Total** |  | **3467** |

**Table 2:** Network inference methods tested in this study (abbreviations in parentheses). Methods can be grouped according to the algorithm family or the regularization type. Algorithm families include correlation, partial correlation with inverse covariance, partial covariance with regression, and mutual information. Regularization types employed by methods can be shrinkage, sparsity, or a combination of shrinkage and sparsity as in ELASTICNET.

| Family | Method |  | Regularization | Running time* (sec) |
| --- | --- | --- | --- | --- |
| Correlation | Pearson correlation (PEARSONCOR) |  | None | 0.022 |
|  | Spearman correlation (SPEARMANCOR) |  | None | 0.053 |
| Partial correlation with inverse covariance | Simple partial correlation (SIMPLEPARCOR) |  | None | 0.060 |
|  | GeneNet shrunken covariance matrix (GENENET) |  | Shrinkage | 0.184 |
|  | Graphical lasso (GLASSO) |  | Sparsity | 0.857 |
| Partial correlation with regression | Partial least squares regression (PLSNET) |  | Shrinkage | 109.370 |
|  | Ridge regression (RIDGENET) |  | Shrinkage | 146.456 |
|  | Lasso regression (LASSONET) |  | Sparsity | 2.99 |
|  | Elastic net regression (ELASTICNET) |  | Sparsity + shrinkage | 6256.607 |
| Mutual information | Algorithm for reconstruction of gene regulatory networks (ARACNE) | additive penalty | None | 3.966 |
|  |  | multiplicative penalty | None | 3.971 |
|  | Context-likelihood of relatedness (CLR) |  | None | 3.995 |
|  | Network inference with maximum relevance / minimum redundancy feature selection (MRNET) |  | None | 4.139 |

\* Running time is for the optimal-parameter runs at limited recall, averaged over 11 tumor types.

The performance comparison for 13 methods was done separately for each tumor type. For a given tumor type, our procedure involved two steps. In the first step, we aimed to put all methods on an equal footing by finding each method’s optimal parameter values. This was achieved by running each method multiple times with different parameter values obtained from a one- or two-dimensional grid, computing the AUPRs for the resulting gene lists, and then finding the parameter or parameter combination with the highest AUPR. The parameters of each method and the design of the grid search are listed in **Table S1**. In the second step, the highest AUPR values from all methods were compared to determine the method with the best performance. This procedure was repeated for each one of the 11 tumor types. Therefore the best-performing method may be different for each one of the tumor types.

There is, however, a caveat concerning the computation of AUPRs from the entire span of the PR curves. We observe in PR curves that (1) there is no significant difference among methods beyond a 10% recall level, and (2) the precision level of network predictions is very low when recall is 10% or higher, suggesting that network predictions are more likely to be affected by noise. The PR curves for BRCA and GBM are shown in **Figure 2a** as representative examples of these two phenomena. The PR curves comparing parameter configurations within each method also exhibited the same pattern (data not shown). Therefore, we chose to use AUPR only from the 0-10% recall range (i.e. **limited-recall**), and not from the entire recall range (i.e. **full-recall**) for the comparison of parameter configurations or the comparison of methods. As the parameter configuration optimizing AUPR in the limited-recall span can be different from that in the full-recall span, some methods were observed to have different PR curves for the limited-recall case (**Figure 2b**). The subsequent analysis is carried out with network predictions from the limited-recall case. The optimal parameter values and the number of edges in the limited-recall case for each method and tumor type are shown in **Table S2** and **Table S3** respectively.

**Figure 2:**
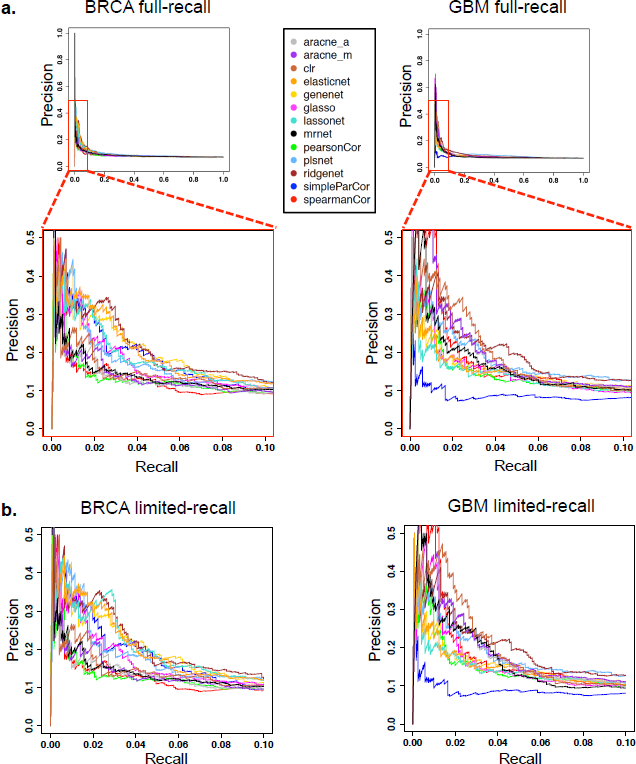
Precision-recall (PR) curves optimized for full versus limited range of recall values. (**a**) ***Top panel:*** PR curves for the 13 methods in the BRCA and GBM cohorts. PR curves are constructed by cumulatively increasing the number of edges from a ranked edge list. For each method, the relevant curve is computed with a choice of parameters that maximize AUPR in the recall range [0,1] (*i.e.* full-recall). ***Bottom panel:*** A zoomed-in version for recall in [0,0.1] and precision in [0,0.5]. (**b**) PR curves when the parameters are chosen to optimize AUPR specifically in the [0,0.1] recall range (*i.e.* limited-recall). We choose the limited-recall case for subsequent analysis because of two reasons. Beyond the 10% recall level, (1) the difference among methods become indiscernible, and (2) the precision level is very low suggesting network predictions are more likely to be affected by noise.

### Performance comparison of network inference methods

After identifying the PR curves to compare the methods, we asked whether any particular method is a clear winner by being the best in all of the 11 tumor types. The AUPR values in **Figure 3a** indicate that there is no single method that performs the best for all investigated tumor types. The tumor types in this figure are ordered from left to right according to increasing coefficient of variation. The differences in the tumor-wise AUPR means and variances indicate that the 11 tumor types are not equally amenable to network inference with RPPA data. These differences could partially be explained by the different statistics of predicted networks such as average-node-degree and network density, which we found to be negatively correlated with AUPR (Spearman r = −0.626 and −0.453 respectively) (**Text S2, Figure S1**).

**Figure 3:**
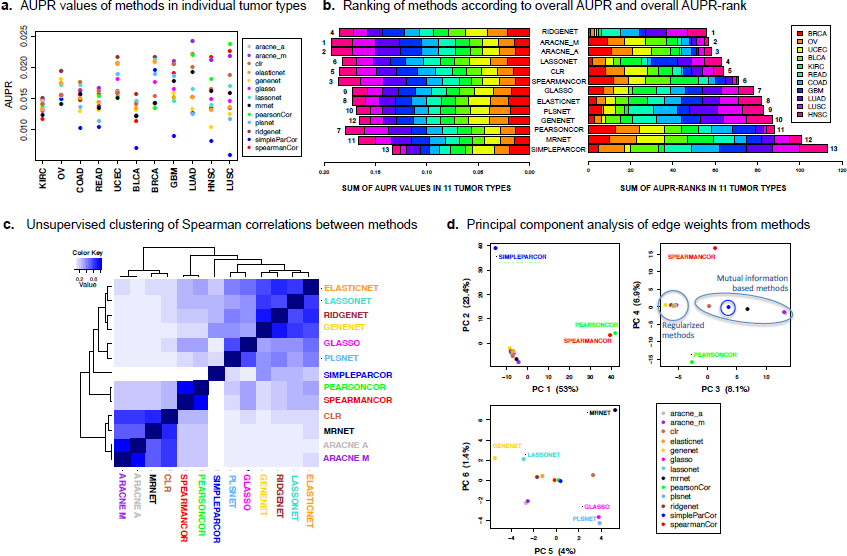
Performance comparison and unsupervised clustering for 13 network inference methods. (**a**) AUPR for each method in individual tumor types. Tumors are ordered according to increasing coefficient of variation. (**b**) Ranking of methods according to (left panel) overall AUPR and (right panel) overall AUPR-rank in 11 tumor types. (**c**) Unsupervised hierarchical clustering of the Spearman correlations between methods. (**d**) Principal component analysis of edge weights from the methods by stacking edge lists from the investigated tumor types.

Given the absence of a clear winner among the methods, we next asked what the overall best-performing methods were. To achieve an overall comparison of the methods, we ranked them across all tumor types based on (1) overall AUPR and (2) overall AUPR-rank. For these two criteria, we computed respectively the sum of a method’s AUPR values in the investigated tumor types (**Figure 3b**, left panel), and the sum of its AUPR ranks in the same tumor types (**Figure 3b**, right panel). The different-colored segments in horizontal bars correspond to tumor types as shown in the legend. The numbers next to the horizontal bars indicate the rank of the method for the relevant criterion. The best rank of 1 is given to the highest overall AUPR but the lowest overall AUPR-rank because higher AUPR values but lower AUPR-ranks indicate better performance.

We observe in **Figure 3b** that the overall AUPR values (left) did not show as wide a variability across methods as the overall AUPR-ranks (right). This might be due to the **overfitting** of the methods to the benchmark network as each method was run with parameters that optimize performance (AUPR) against the same benchmark. The small differences in overall AUPR values suggest that these methods may have a general capacity to achieve similar performance in other contexts as long as their respective parameter space is sufficiently explored. However, such similarity in performance does not preclude the possibility that some methods consistently outperform others even if by small margins. To investigate this possibility, we ordered the methods from top to bottom according to increasing overall AUPR-rank. This choice in the ordering shows that *RIDGENET* is the best-performing method overall. Broken down by tumor type, *RIDGENET* is the best for BRCA, OV, UCEC, BLCA and KIRC; but is not as good as *ARACNE* variants for HNSC, LUSC, LUAD, GBM, COAD, and READ. On the poor performance side, *SIMPLEPARCOR* has the worst rank according to both the overall AUPR and the overall AUPR-rank (**Figure 3b**).

### Network predictions cluster methods primarily based on algorithm family

We next investigated the level of similarity among the network predictions of all 13 methods. One question here is whether the network predictions, as given by the inferred edge weights, would cluster the methods according to shared properties such as the regularization technique or the algorithm family listed in **Table 2**. To this end, we created one vector for each method by stacking the relevant edge weights from all 11 tumor types. We then computed the *Spearman correlation* between each pair of the methods, and also performed dimensionality reduction on the same vectors using *principal component analysis* (PCA). Unsupervised clustering on the Spearman correlation matrix (hieararchical clustering with complete linkage and Euclidean distance) and PCA on the edge weight matrix reveal concordant results in terms of the grouping of the methods (**Figure 3c–3d**). We observe three major groups of methods in **Figure 3c**: (1) Mutual-information-based methods *ARACNE* (variants)*, CLR, MRNET*, (2) correlation-based methods *SPEARMANCOR* and *PEARSONCOR*, and (3) partial-correlation-based methods. *SIMPLEPARCOR* from the third group can be considered an outlier compared with the other partial-correlation methods. Therefore, if we remove it as a separate group, the remaining partial-correlation methods *RIDGENET, LASSONET, ELASTICNET, PLSNET, GLASSO, GENENET* can also be categorized as **‘regularized methods’**.

In the PCA plots, the 1^st^ princial component (PC) primarily separates correlation-based methods *SPEARMANCOR* and *PEARSONCOR* from the others, accounting for 53% of the variance (**Figure 3d**). Correlation methods are fundamentally different from other investigated methods because they do not attempt to eliminate transitive edges in any way. This defect could predict poor performance for both *SPEARMANCOR* and *PEARSONCOR*. However, the fact that the rank-based *SPEARMANCOR* achieves a superior overall performance compared to the value-based *PEARSONCOR* (**Figure 3b**) suggests that outliers in the data can bias the Pearson correlation strongly enough to substantially reduce the quality of the predicted network.

The 2^nd^ PC (23.4% variance) separates *SIMPLEPARCOR*, a method that is based on Gaussian graphical models and that employs the sub-optimal pseudo-inverse technique when the covariance matrix is singular. Even when the covariance matrix is non-singular, the inversion of the covariance matrix without any regularization is known to introduce defects into the inference procedure unless the number of samples is at least twice the number of features ^16^. As the cohort sizes in this study are less than twice the number of proteins (2*187=374) for 7 of the 11 tumor types (**Table 1**), it is not surprising that *SIMPLEPARCOR* has poor performance in these tumor types, hence the poorest overall performance by a margin (**Figure 3b**). Indeed, we can observe that the tumor types where *SIMPLEPARCOR* achieves relatively better ranks are BRCA, OVCA, KIRC, and UCEC, the 4 tumor types that have cohort size greater than 374 (**Figure 3a–3b** and **Table 1**).

The 3^rd^ PC (8.1 % variance) achieves the separation of mutual-information methods from regularized methods. Mutual-information-based methods have the capability to model nonlinear relationships, but are not able to infer the direction of the relationship. These two fundamental differences may account for the clear separation of these methods from the others. Principal components can achieve a separation of regularization-based methods only at the 5^th^ and 6^th^ PC, which account for as little as 4% and 1.4% of the variance respectively (**Figure 3d**).

### TOP6: a group of high-performers instead of a *“best”* method

The modest differences between overall AUPR values in the left panel of **Figure 3b**, and also the lack of a consistently best-performing method in all tumor types are reasons to refrain from recommending one method as the best off-the-shelf method for PPI inference. Therefore, we propose a set of **high-performers** by taking into consideration both the overall AUPR and the overall AUPR-rank criteria. The methods that rank in the top six according to both of these criteria are the same six methods: *RIDGENET, ARACNE-M, ARACNE-A, LASSONET, CLR,* and *SPEARMANCOR* (**Figure 3b**). This set of high-performers, referred to as **TOP6** from here on, includes representative methods from all algorithm families in **Table 2** except for inverse-covariance-based partial-correlation methods. This may be indicative of inverse covariance being a poor framework to model PPIs in cancer if especially the cohort size is not several times as large as the number of proteins. In conrast, linear measures such as correlation and (*ℓ*_1_- or *ℓ*_2_-regularized) partial-correlation, and also nonlinear measures such as mutual-information are all represented in the set of high-performers.

### Determining the *“consensus”* edges for the unsupervised clustering of tumor types

We next asked how the network predictions from the TOP6 methods cluster the 11 tumor types. However, similar to the reduction from 13 methods to the TOP6 methods, it was necessary to apply a significance threshold for edges before performing the clustering. P-values were not a viable option as significance scores, because several methods did not return p-values. Even if p-values were obtained from all methods, it would not be possible to combine the p-values in this study in a statistically sound way because all methods used the same data, hence violating the independence requirement. Therefore, we resorted to an alternative method to obtain ***consensus*** significance scores for edges.

We computed, for a given tumor type, (1) *consensus edge ranks* by taking the average of ranks from the TOP6 methods, and (2) *consensus edge weights* by taking the average of weights again from the TOP6 methods. The consensus ranks formed the basis for our significance levels, while the consensus weights were used in the clustering steps. Comparing consensus-edge-ranks obtained from the TOP6 methods with those obtained from all 13 methods (ALL13) showed that the TOP6 methods yielded slightly higher AUPR than ALL13 against the PC gold-standard (**Figure S3b, Text S3)**. This finding confirmed the use of TOP6 as a superior choice over ALL13.

The number of edges to use for the unsupervised clustering of tumor types was determined in the following way. For a certain threshold, we extracted all edges from a given tumor type that have a consensus-edge-rank smaller (more significant) than the threshold level. We then formed a matrix of edges by tumor types by combining extracted edges from all 11 tumor types. Next, we computed the principal components (PCs) constructed as a linear combination of the tumor-type-vectors, and inspected the percentage of variance explained by the first three PCs. By increasing the threshold from 1 to 2000, we observed that using a consensus-rank threshold of 425 allowed an optimal separation of tumor types along the first three PCs (**Figure S4, Text S4).**

### Unsupervised clustering of tumor types by edge weights recapitulate recently published gene- and protein-expression-based groups

Using the consensus-rank threshold of 425, we investigated the natural groupings in the set of 11 tumor types when each tumor type was represented with the consensus-edge-weights obtained from the TOP6 methods. The number of edges in each tumor type that pass the consensus-rank threshold is shown in **Table S4**. The union set of these significant edges from the tested tumor types has 1008 edges. We refer to this union set as the ***discovery set***, and use it to perform principal component (PC) analysis and hierarchical clustering of tumor types.

We see in the PC analysis that PC1 and PC2 jointly separate the 11 tumor types into three groups, and also that PC3 further breaks down one group into two to result in a total of ***four groups***: 1) COAD, READ; 2) LUSC, LUAD, HNSC; 3) GBM, KIRC; and 4) OV, BRCA, BLCA, and UCEC (**Figure 4a**). These results are concordant with the clusters from hierarchical clustering (**Figure 4b** dendrogram) and also with the previously defined Pan-Cancer groups in the literature as we elaborate below.

**Figure 4:**
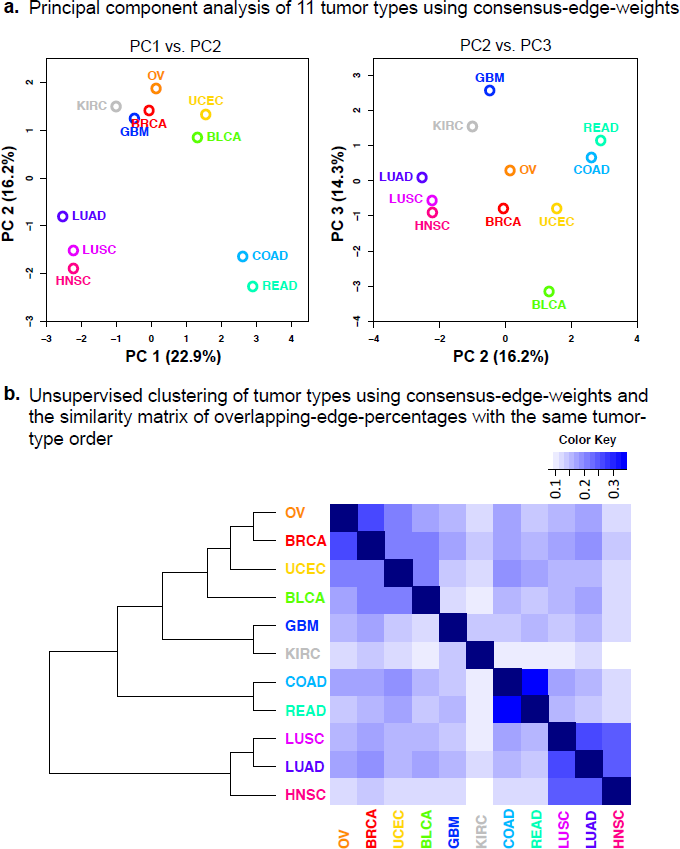
Principal component analysis and unsupervised clustering of 11 tumor types using consensus edge-weights from the TOP6 methods. (**a**) Four major groups of tumor types can be observed in the PC1 vs. PC2 (left) and PC2 vs. PC3 (right) plots: 1) COAD, READ; 2) LUAD, LUSC, HNSC; 3) GBM, KIRC; 4) OV, BRCA, UCEC, BLCA. (**b**) Hierarchical clustering (Ward linkage and Euclidean distance) on consensus-edge-weights places tumor types into the same four groups on the dendrogram (left). The heat-map on the right is constructed from the percentages of overlapping edges between tumor types. The order of tumor types in the heat-map is taken from the dendrogram on the left.

As for the first group, COAD and READ have previously been shown to cluster together in the Pan-Cancer subtypes defined both by RNA expression (k=13)^27^ and by protein expression (k=8)^4^. These tumors have also been shown to have common DNA-based drivers (mutations and somatic copy number alterations), and hence been treated as one disease ^2, 3, 28^. Our finding that COAD and READ have the highest percentage of shared PPIs in this study (**Figure 4b** heat-map) is also in line with these observations. Note that the order of tumor types in the heat-map is taken from the dendrogram on the left, and that each cell represents the fraction of the intersection set over the union set of edges from two tumor types.

The tumors in the second group (LUSC, LUAD, and HNSC) have also been previously assigned to a single Pan-Cancer subtype in terms of protein expression ^4^. However, RNA expression and somatic copy-number alteration (SCNA) data types have divided these tumor types into two groups: (1) a squamous-like subtype including HNSC and LUSC, and (2) a separate LUAD-enriched group ^3, 27^. In contrast to this separation where cell histology plays a more important role, both protein expression levels and PPI weights primarily separate these three tumor types based on tissue of origin: (1) Lung-derived tumors LUAD and LUSC, and (2) a separate HNSC group (**Figure 4b** dendrogram and ^4^).

Tumors in the third and fourth groups (GBM, KIRC, OV, UCEC, BRCA, and BLCA) can be separated along a continuum in the PC3 dimension (**Figure 4a**). However, we can consider GBM and KIRC as a separate group as these two tumor types separate from the other four in the unsupervised clustering dendrogram in **Figure 4b**. GBM and KIRC also cluster most closely among this set of 11 tumor types according to somatic copy-number alterations and protein expression levels ^3, 4^. However, KIRC also shows an outlier behavior for PPI networks in that it exhibits the lowest fraction of shared PPIs with other tumor types (**Figure 4b**). GBM, on the other hand, has an outlier property by being on one extreme of the separation along the PC3 dimension. This may reflect the fact that GBM samples arise from **glial cells** in the brain, a histological origin that shows marked differences from epithelial cells. Indeed, GBM was previously shown to have a homogeneous cluster comprised of only GBM samples in terms of both RNA and protein expression levels ^4, 27^.

The fourth group contains OV, UCEC, BRCA, and BLCA; the first three of which can be categorized as women’s cancers. The proximity of women’s cancers in clustering results may point to female hormones, such as estrogen and progesterone, causing a similar profile of PPI weights. BLCA is most similar to women’s cancers (**Figure 4b**), but it also is on one extreme of the separation along the PC3 dimension. This is concordant with the previously discovered Pan-Cancer subtypes because BLCA was shown to have the characteristic property of being one of the most diverse tumor types in the TCGA Pan-Cancer dataset. It had samples in 7 major RNA expression subtypes, and histologies in squamous, adenocarcinoma, and other variants in bladder carcinoma ^27^. Next, we performed unsupervised clustering on the 1008 edges in the *discovery set* to investigate edges that are specific for a certain tumor type and those that are shared between two or more tumor types.

### Unsupervised clustering of edges reveals two modules of positive interactions highly recurrent across the tested tumor types

An unsupervised inspection of the 1008 PPIs in the discovery set shows that these edges form three main groups: (1) a large group of positive edges with low or no recurrence in tumor types (793 edges), (2) a small group of positive edges with high recurrence (123 edges), and (3) a small group of negative edges with low or no recurrence (92 edges) (**Figure 5**). In this set of most significant edges, both the number and the overall weight of negative edges are smaller with respect to positive edges. This may indicate either the lower prevalence of mutual exclusivity relationships for *in vivo* protein-protein interactions, or merely the difficulty of discovering negative PPIs from RPPA data.

**Figure 5:**
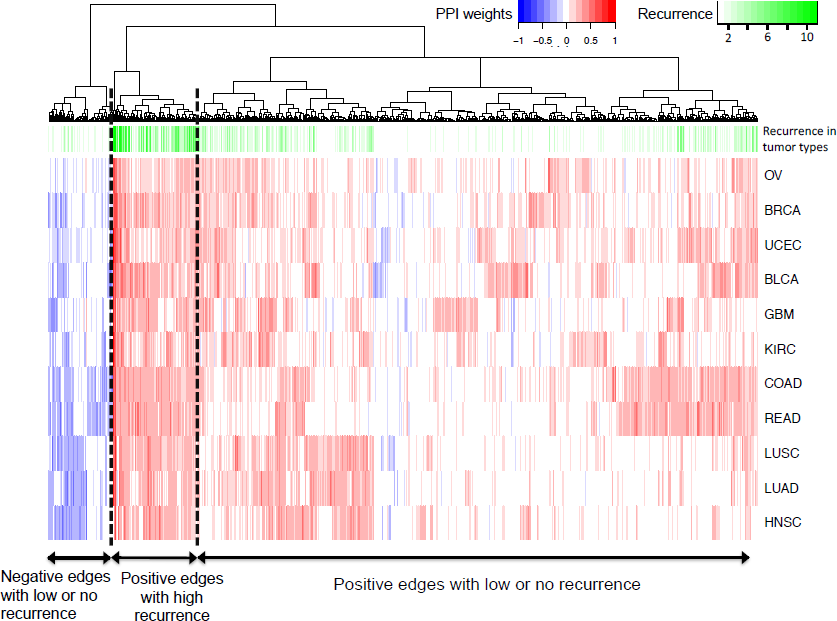
Hierarchical clustering of the 1008 edges in the *discovery set* (Ward linkage and Euclidean distance). Consensus edge-weights are plotted in the heat-map with **blue** denoting negative, and **red** denoting positive edges. Recurrence of edges in tumor types ranges from 1 to 11, and is denoted with shades of **green**. Three edge groups can be observed on the dendrogram: (1) positive edges with low or no recurrence in tumor types, (2) positive edges with high recurrence, and (3) negative edges with low or no recurrence.

Next, we visualize the highly recurrent positive edges (group 2) on a network layout to investigate their biological significance (**Figure 6**). It is remarkable that the network of positive-higly-recurrent interactions can be separated into two distinct modules: one including signaling events (interactions that involve at least one phosphoprotein) and the other including only non-signaling interactions without any phosphoproteins. The existence of an exclusively signaling module in the set of highly recurrent positive interactions underlines the ***pan-cancer importance of signal transduction events***, particularly the RAS/MAPK pathway and the PI3K/AKT/mTOR pathway (magenta and red nodes in **Figure 6** respectively). The interaction partners for the group 1 and group 3 edges are given in **Supplementary File XX.**

**Figure 6:**
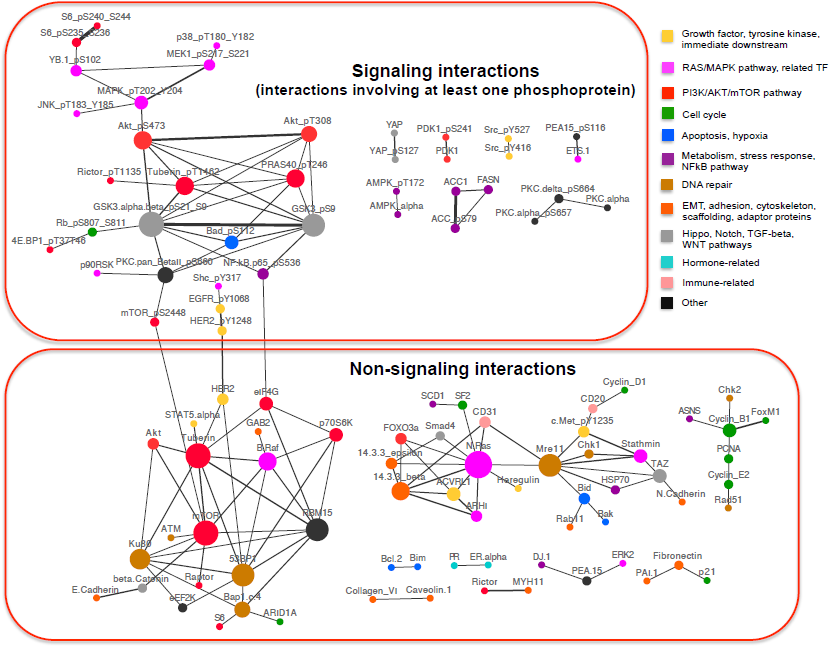
A network view of the positive highly recurrent edges. Signaling interactions (interactions involving at least one phosphoprotein) and non-signaling interactions (interactions that do not involve any phosphoproteins) can be placed in two separate modules that are connected with only a few interactions involving the mTOR, HER2, and NFκB signaling activities. Node size denotes degree (number of edges incident on the node), and node color denotes biological function as shown in the legend. The edge widths are drawn proportionally to the consensus edge weights.

The information flow between these two communities can go through one of two major interactions; one between total mTOR and phospho-mTOR, and another one between phospho-NFkB and eIF4G. The interaction between HER2 and phospho-HER2 cannot be used for information flow because it does not connect the two main bodies of the signaling and non-signaling interactions. Moreover, the link between phospho-HER2 and phospho-EGFR is most likely an artifact due to antibody specificity problems. We also observe in **Figure 6** that the strongest interactions inferred from the PANCAN11 RPPA data are those between two different phosphorylation states, or between the phosphorylated and unphosphorylated states, of the same protein. The examples to the former are (1) phospho-S6, (2) phospho-Akt, and (3) phospho-GSK3 interactions. The example to the latter is the interaction between ACC1 and phospho-ACC.

### Over-representation of *discovery* set edges in REACTOME & KEGG gene lists as a means of *interpretation*

The network visualization of the discovery set PPIs presents an opportunity to *discover* biologically interesting cancer-related interactions. However, when the number of edges is too large such as the group 1 low-or-no-recurrence positive edges, the high inter-connectedness of proteins makes it hard to *interpret* network results. Thus, in order to to facilitate the *interpretation* of the discovery set edges, we sought to identify the REACTOME ^21^ and KEGG ^23^ gene lists in which these edges are overrepresented via a PPI-enrichment analysis.

Gene/protein set enrichment tools exist in abundance in the literature, however are not applicable for PPI enrichment because PPIs have to be compared with an interaction knowledgebase such as Pathway Commons. Therefore, we designed a custom PPI enrichment procedure that achieves the translation from a PPI-by-tumor-type matrix (such as the discovery set) to a gene-list-by-tumor-type matrix via an intermediary PPI-by-gene-list matrix (**Online Methods** and **Figure S5**).

The PPI enrichment analysis shows both gene lists that are recurrent in multiple tumor types and also gene lists that are specific for one or two tumor types. Similar to the group 2 (highly recurrent positive) edges that showed an important signaling component in **Figure 6**, we observe in **Figure 7** that gene lists with the strongest enrichment results primarily contain **signal transduction** events such as receptor tyrosine kinase signaling (EGFR, ERBB2, ERBB4, FGFR, KIT, VEGFR, PDGFR), RAS/MAPK signaling, PI3K/AKT/mTOR, and HIF signaling. The other major group of gene lists with the strongest enrichment concerns **innate and adaptive immunity** related interactions. Some examples are signaling interactions involving DAP12, B-cell receptor, T-cell receptor, Toll-like receptor, and TNF. Several gene lists such as Hippo signaling and Wnt signaling had strong enrichment results for only a subset of the tumor types.

**Figure 7:**
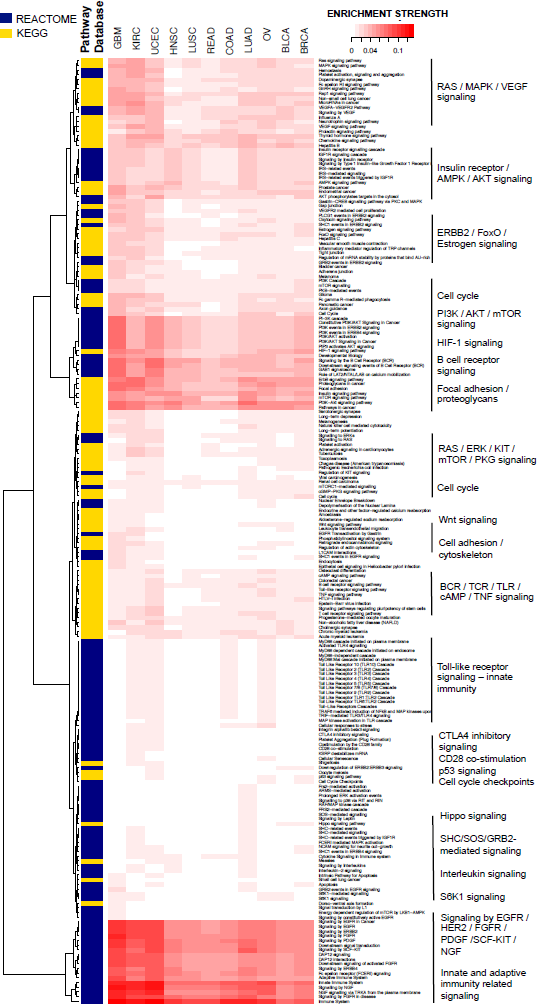
PPI enrichment in REACTOME and KEGG gene lists. recapitulates the pan-cancer importance of signal transduction events, and further underlines the centrality of innate and adaptive immunity related signaling in tumor biology.

## DISCUSSION

Discovering protein-protein interactions in cancerous cells is an important but challenging goal. In this study, we computationally inferred proteomic networks in 11 human cancers using 13 different methods, and presented a performance comparison of the methods accepting a simplified reference network from the Pathway Commons (PC) information resource, which is based on experiments and publication digests, as the standard of truth. PC is a collection of curated interactions from many different normal and disease conditions (a formal and computable representation of available pathways and interactions). We acknowledge that a complete standard of truth for pathways is not currently available and that our methodology is therefore subject to certain caveats as discussed below. Despite these caveats, computational inference of protein-protein interaction networks from measurements of protein levels across a set of conditions are attractive in that they can reduce the hypothesis space of interactions and inform researchers on the potentially active pathways in the experimental model.

Our comparison of the performance of network inference methods indicates that no single method has the best performance in all tumor types, but a group of six methods, including diverse techniques such as correlation, mutual information, and regression, consistently rank highly among the tested methods. These six methods consist of *RIDGENET* and *LASSONET,* ridge and lasso regression based partial correlation methods employing an *ℓ*_2_ and *ℓ*_1_ penalty respectively; *ARACNE-A*, *ARACNE-M,* and *CLR,* mutual-information methods that differ based on their penalty type or the choice of standardization for mutual information; and SPEARMAN CORRELATION, which assesses the strength of the linear relationship between the *ranks* of the values in two same-length vectors. From a tumor type perspective, we find that not all tumor types are equally amenable to network discovery with RPPA data. Five tumor types (KIRC, OV, COAD, READ, and BLCA) consistently had lower-AUPR predictions by all network inference methods.

A consensus network from the group of high-performing methods reveals that the strongest protein-protein interactions that are shared across the tested tumor types are receptor tyrosine kinase (RTK)-related and immunity-related signaling pathways. Other strong interactions that are shared across most tumor types include the RAS/MAPK, PI3K/AKT/mTOR, HIF signaling pathways as well as immunity-related pathways such as B-cell receptor, T-cell receptor, Toll-like receptor, and TNF signaling.

The caveats in our workflow as shown in **Figure 1** concern both the *computational inference* and the *pathway knowledgebase* arms of the analysis. In the computational inference arm (Steps 1 and 2), the caveats include questions around (1) the quality of RPPA experiments and whether the signal-to-noise ratio in RPPA experiments is high enough to allow the inference of direct interactions, and (2) the reliability of results from computational network inference methods (**Text S1**). In the pathway knowledgebase arm (Steps 3-5), the fidelity of pathway models in knowledgebases is limited due to factors including (1) the quality of wet-lab experiments for PPIs such as yeast-2-hybrid ^29^, (2) missing or inaccurate information in the database due to poor curation, (3) the lack of context information for PPIs, such as cell or tissue type or physiological conditions, and (4) the ascertainment bias in the knowledgebase (primarily incomplete coverage) as discussed in the **Introduction**. More generally, pathways in knowledgebases such as Pathway Commons are only model descriptions of reality typically summarizing a set of experiments and do not represent an absolutely ‘true’ (and certainly not complete) set of interactions.

In a recent multi-method comparison study for gene network inference, regression-based methods were represented mostly by modifications of the *ℓ*_1_-penalized lasso algorithm; however methods involving an *ℓ*_2_ penalty, such as ridge regression or elastic net, were not included ^30^. Moreover, the *ℓ*_1_-penalized methods did not achieve the best-overall performance in gene network inference. We find in this study that *ℓ*_2_-penalized methods such as ridge regression can outperform the lasso in the inference of proteomic networks. Even though the concurrent execution of feature selection and model-fitting may appear to be an attractive property for lasso-regression, we recommend performing an unbiased test for both *ℓ*_1_ and *ℓ*_2_-penalized models in the exploratory phase of a study. It is not guaranteed that the variables selected by the *ℓ*_1_ penalty will be the most biologically important ones in the system.

In future work, it will be important to assess the predictive power of the inferred PPI networks. For example, it would be useful to evaluate these networks in terms of how much they assist in the understanding of oncogenesis, response to therapy, and design of combination therapies that deal with feedback loops. It is also desirable to incorporate time-dependent readouts from perturbation experiments to be able to build causal models and enhance the predictive power of proteomic networks. An obstacle against building causal models, such as Bayesian networks, with the PANCAN11 RPPA data, was the relatively large size of the network (187 nodes) compared with the number of available samples in individual tumor types (between 127 and 747). Probabilistic models such as Bayesian networks require at least an order of magnitude larger number of samples for a sound estimation of model parameters.

The significance of this work extends beyond cancer. Discovering direct, potentially causal interactions between proteins is an opportunity in all areas of molecular biology where proteins are measured in different conditions, and where correlations are informative. The methodology presented here can easily be adopted to study interactions in different molecular biology contexts.

## ONLINE METHODS

### Dataset

The pan-cancer reverse phase protein array (RPPA) dataset was downloaded from ***The Cancer Proteome Atlas***^5^ on April 12, 2013. This dataset is denoted as PanCan11 and contains protein expression data for 187 proteins and 3467 tumor samples. The 11 tumor types represented in this dataset are bladder urothelial carcinoma (BLCA), breast invasive carcinoma (BRCA), colon adenocarcinoma (COAD), glioblastoma multiforme (GBM), head and neck squamous cell carcinoma (HNSC), kidney renal clear cell carcinoma (KIRC), lung adenocarcinoma (LUAD), lung squamous cell carcinoma (LUSC), ovarian serous cystadenocarcinoma (OVCA), rectum adenocarcinoma (READ), and uterine corpus endometrioid carcinoma (UCEC).

PanCan11 patient samples were profiled with RPPA in different batches, and normalized with replicate-based normalization (RBN). RBN uses replicate samples that are common between batches to adjust antibody means and standard deviations so that the means and standard deviations of the replicates become the same across batches.

### Pathway Commons query with PERA

Pathway Commons (PC) stores pathway information in BioPAX ^22^ models that contain formal computable representations of diverse events such as biochemical reactions, complex assembly, transport, catalysis, and physical interactions. We queried PC with the “prior extraction and reduction algorithm” (PERA)^26^ for the proteins and phosphoproteins in the PANCAN11 RPPA dataset.

PERA is a software tool and a protocol that, given a set of observable (phospho- and/or total) proteins, extracts the direct and indirect relationships between these observables from BioPAX formatted pathway models ^26^. PERA accepts a list of (phospho)proteins identified by their HGNC symbols, phosphorylation sites and their molecular status (active, inactive or concentration/total protein) as input and based on the pathway information provided by the Pathway Commons information resource ^20^, it produces a binary and directed network. The biggest advantage of PERA over other similar tools, such as STRING ^31^ or GeneMania ^32^, is that it considers not only the name/symbol of a protein but also its phosphorylation states – enabling finer mapping of entities and pathways.

### Implementing network inference methods in R

The analysis was performed using the R language ^33^. The R functions used to implement the network inference methods are as follows: The `cor` function in the **stats**^34^ package for PEARSONCOR and SPEARMANCOR; the `ggm.estimate.pcor` and `cor2pcor` functions in the **GeneNet**^35^ package for GENENET and SIMPLEPARCOR; the `ridge.net, pls.net`, and `adalasso.net` functions in the **parcor**^36^ package for RIDGENET, PLSNET, and LASSONET; the g`lasso` function in the **glasso**^37^ package for GLASSO; the `aracne.a, aracne.m, clr`, and `mrnet` functions in the **parmigene** package ^38^ for ARACNE-A, ARACNE-M, CLR and MRNET. The ELASTICNET method was implemented as a modification of the `adalasso.net` function in the **parcor** package, and is available upon request. Mathematical descriptions of the algorithms used are provided in **Text S6**.

### Evaluating performance of network inference methods

As all of the algorithms we studied in this work provided undirected network predictions, we converted the PERA output to undirected edges to arrive at the benchmark edge list (‘benchmark network’) used in this study. We then constructed a series of precision-recall (PR) curves for each algorithm interrogating their performance with a range of values for their respective parameters (**Table S1**). *Precision*, is the fraction of the number of correctly predicted edges (predicted edges that can be found in PC) to the number of all predicted edges. *Recall*, on the other hand, is the fraction of correctly predicted edges to the number of all edges in PC.

The PR curve for a given parameter configuration was constructed by taking the edge list ranked from the most significant to the least, and then iterating over the edges so that we obtained, at each iteration, a cumulative edge set that included all the edges seen up to and including that iteration. For each iteration, we computed the precision-recall value pair for the edge set and placed this value pair on the PR plot. We plotted a separate PR curve for each parameter configuration for the nine methods that required specification of parameter values (all methods except *PEARSONCOR, SPEARMANCOR, SIMPLEPARCOR,* and *GENENET*). The PR curve that had the greatest area under the curve (AUPR) between the [0,0.1] recall range (i.e. limited-recall) was identified as the optimal PR curve for that particular method. The optimal parameter values for the limited-recall case are shown in **Table S2**. For methods that did not have user-specified parameters, there was only one PR curve and that was adopted as the optimal PR curve. In the subsequent step, the AUPRs from the optimal PR curves were compared to be able to rank the methods and evaluate their performance relative to the benchmark network.

The steps involved in computational network prediction and performance evaluation are discussed in detail in **Text S5**.

### Rationale to prefer high precision over high recall

We find that the inferred interactions in various tumor types are a relatively small subset of the benchmark network derived from PC (*i.e.* low recall). Low levels of recall are readily acceptable for satisfactory performance because it is expected that interactions inferred from a single disease (cancer) and a single cancer type will not retrieve all of the interactions in the PC benchmark. However, it is desirable that, when an algorithm calls an interaction, there is a high probability that this inference is correct, *i.e.* high levels of precision are essential for nominating a network inference method as competitive.

### PPI enrichment in REACTOME and KEGG gene lists

The 187 antibodies in our RPPA dataset correspond to 151 unique genes. The outline of the steps involved in PPI enrichment analysis is as follows: (1) We provided the 151-gene set as input to ClueGO^39^ to inspect enrichment of the genes in REACTOME^21^ and KEGG^23^ gene lists. The output gene lists with significant enrichment function as a universal set of all possible gene lists in which PPIs of the same set of 151 genes can be enriched in. We obtained a binary matrix of genes versus gene lists from this step.

Next, (2) a filtering step was performed separately for each tumor type to convert the “gene by genelist” matrix to a PPI-by-gene list matrix. A PPI in the discovery set was declared to be enriched in a gene list if both interaction partners (genes as opposed to proteins) were found to be enriched in that gene list in the gene-by-gene list matrix. (3) The binary “PPI by gene list” matrix formed this way was then adjusted for the strength and sign of the PPIs to allow for a weighted voting scheme. (4) This weighted matrix then allowed the computation of tumor-specific gene list enrichment scores by simply taking the average of weights for each gene list across the interactions. (5) The gene list enrichment scores for each tumor were then combined in a separate matrix to form the “gene list by tumor type” matrix. To achieve better visualization in hierarchical clustering, we filter out the gene lists in this matrix that have weights less than 0.01 for all tumor types.

## ACKNOWLEDGEMENTS

The authors thank Ö. Babür, E. Demir, B.A. Aksoy, E. Reznik, and A. Korkut for their helpful discussion on the manuscript.

